# Testosterone eliminates strategic prosocial behavior through impacting choice consistency in healthy males

**DOI:** 10.1101/2022.04.27.489681

**Authors:** Hana H. Kutlikova, Lei Zhang, Christoph Eisenegger, Jack van Honk, Claus Lamm

## Abstract

Humans are strategically more prosocial when their actions are being watched by others than when they act alone. Using a psychopharmacogenetic approach, we investigated the endocrinological and computational mechanisms of such audience-driven prosociality. 192 male participants received either a single dose of testosterone (150 mg) or a placebo and performed a prosocial and self-benefitting reinforcement learning task. Crucially, the task was performed either in private or when being watched. Rival theories suggest that the hormone might either diminish or strengthen audience-dependent prosociality. We show that exogenous testosterone fully eliminated strategic, i.e., feigned, prosociality and thus decreased submission to audience expectations. We next performed reinforcement-learning drift-diffusion computational modeling to elucidate which latent aspects of decision-making testosterone acted on. The modeling revealed that testosterone compared to placebo did not deteriorate reinforcement learning per se. Rather, when being watched, the hormone altered the degree to which the learned information on choice value translated to action selection. Taken together, our study provides novel evidence of testosterone’s effects on implicit reward processing, through which it counteracts conformity and deceptive reputation strategies.

## Introduction

Humans behave more prosocially when their actions are watched by others [1]. This phenomenon has been demonstrated across a variety of social behaviors, such as blood donations [2], church offerings [3], or monetary donations to charitable organizations [4], and is often referred to as strategic prosociality [5], or the audience effect [6]. From an evolutionary perspective, making one’s generosity visible to others has an important signaling value, in that it advertises an individual’s qualities as a potential partner or a valuable group member [7]. In the present study, we propose and investigate whether the steroid hormone testosterone plays a crucial role in shaping such audience effects.

Research in the past decade has demonstrated that testosterone is implicated in a wide spectrum of socially dominant behaviors [8,9]. Exogenous testosterone alleviates subordination to the dominance of others [10-12] and reduces the physiological response to being evaluated by others [13]. Given that enhanced submission to audience expectations has been associated with intense apprehension about social evaluation [14], one possible prediction is that testosterone administration will decrease audience effects.

Contrasting with this view, the hypothesis that testosterone drives status-seeking via reputation building rather than dominance [15, 16] would predict that based on the social context, testosterone might conditionally promote prosocial and especially socially desirable behavior to build up a reputation and increase status.

The present paper is the first that aimed to distinguish between these two alternatives of boosting one’s social status that testosterone may act on. One option is that, in line with the social dominance hypothesis [17], the hormone prioritizes dominant status-seeking and would hence diminish the submission to audience expectations. The other option is that testosterone primarily promotes reputable status-seeking [15, 16]. If true, the hormone could increase strategic prosocial behavior.

Through what neurobiological pathways could testosterone modulate such complex social behaviors? Previously, exogenous testosterone was found to increase dopamine levels in the rat ventral striatum [18], suggesting that the hormone exerts its effects through the modulation of dopaminergic activity in reward-related neural circuits. Besides this insight from animal research, testosterone and reward processing have also been linked in humans [19, 20]. It remains to be shown, though, which specific aspects of reward processing testosterone acts on. For one, during value learning, testosterone may influence the incorporation of so-called prediction errors (PE), which track the difference between predicted and actual outcome [21] and are encoded by the phasic activity of midbrain dopaminergic neurons projecting to the ventral striatum [22, 23]. Alternatively, testosterone may impact the conversion of the learned values into action selection, or the temporal dynamics of the evidence accumulation.

The present study thus not only aimed to investigate if testosterone influences strategic prosociality, but also whether this is achieved by impacting reward-related computations. We employed a novel modeling approach, by combining reinforcement learning with diffusion decision models (RLDDM). This provided a more comprehensive account of the latent processes involved in prosocial decision-making than previous separate RL and DDM approaches [21,24]. Besides describing how subjective values of the choice options are learned through PEs (*learning rate parameters*) and converted to actions (*choice consistency parameter*), the new combination of reinforcement learning and diffusion decision modeling also enabled us to explore the temporal dynamics of these latent processes (*decision threshold and drift-scaling parameters,* see *SM Table S4* for parameter description) [24].

Male participants underwent a double-blind, between-subject, placebo-controlled, testosterone administration and then performed a reinforcement learning task (Figure 1). To compare self- and other-oriented decision-making, participants completed the task for themselves and for an NGO of their choice. While charitable donation tasks [16] and neuroeconomic games [8, 9] classically measure participants’ overall prosociality using deliberated decisions, such as deciding how much money to share with another person, the RL task allowed us to furthermore characterize the hidden individual steps in the process of learning about the consequences actions have for oneself and others (see [25, 26] for similar recent approaches). Critically, the task was performed either in private or when being watched (see *Materials and Methods*).

**Figure 1.**
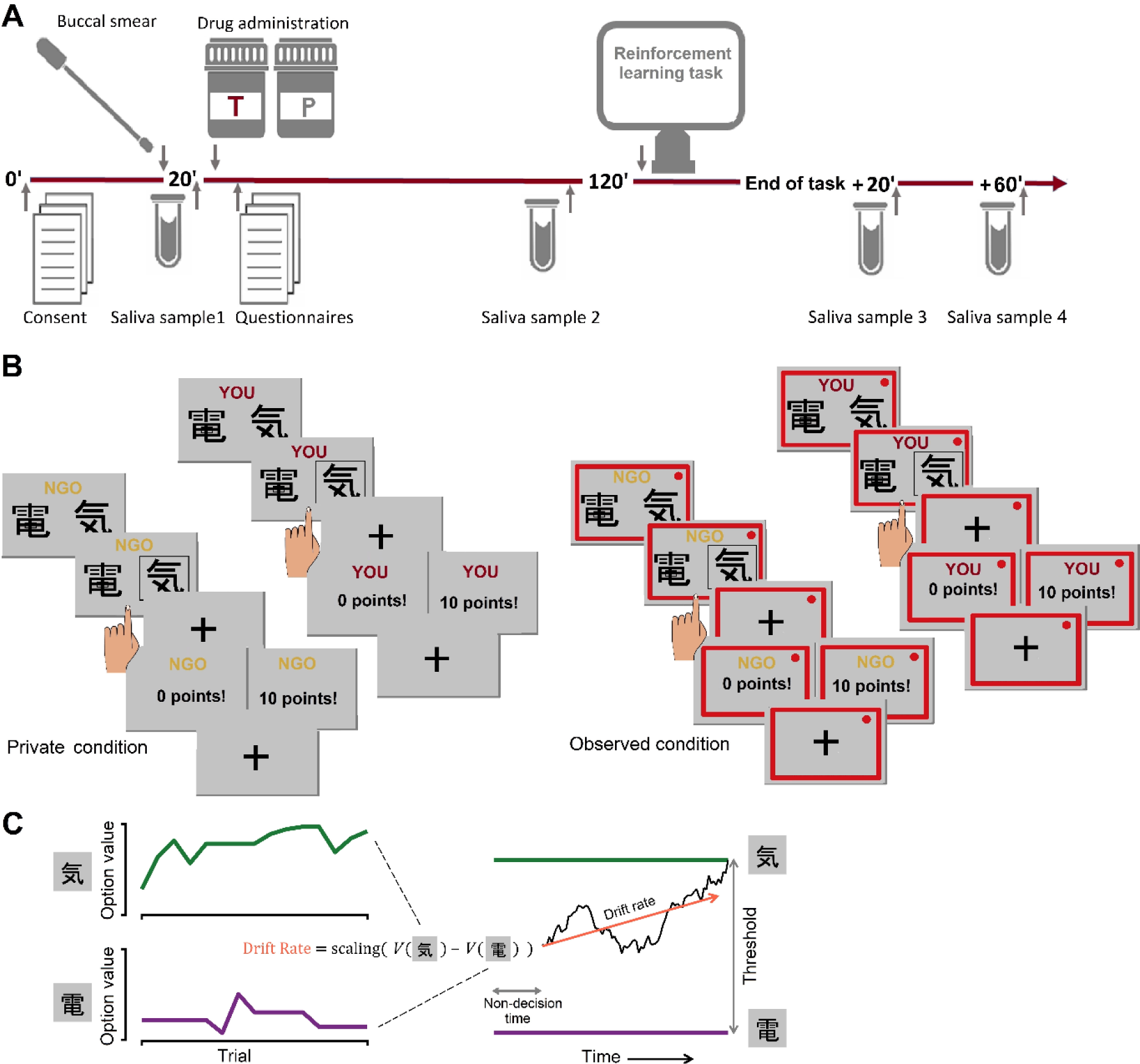
Experimental design and task. *(A)* Timeline of the experimental session. *(B)* Prosocial reinforcement learning task. Participants performed the task either in private or watched by an observer introduced as an NGO association representative. The observation was signaled by a red frame. Each participant completed three blocks of 16 trials for themself and three blocks of 16 trials to benefit an NGO of his choice. *(C)* Schematic of the reinforcement learning drift diffusion model (RLDDM). Left panel: trial-by-trial value updates in RL; right panel: evidence accumulation in DDM. Importantly, the drift rate in DDM is calculated from the value difference between choice options in RL.

Based on previous audience-effect research [2-6], we predicted that when the participants are watched, they will be relatively more prosocial (i.e., make more correct choices for the other vs self) than in private. Crucially, we expected that such an audience effect will be underpinned by relatively faster incorporation of the PEs (captured by the learning rate parameter α in RL); higher consistency in converting values to action probability (captured by the inverse temperature parameter tau in RL, also known as value sensitivity, exploration parameter, or 1/β); and more integrated evidence necessary for making a decision (captured by the threshold parameter in DDM). In other words, participants would learn more efficiently, learned values would inform their behavior more consistently, and their decisions would be more cautious.

Our main hypothesis was that the effects of being watched on other-vs. self-benefitting behavior will be modulated by testosterone administration. Given that testosterone reduces submission signals and stress responses to the social evaluation, allowing for dominant status-seeking [10-13], we hypothesized that testosterone would reduce the audience effect expected in the placebo group. As an alternative prediction, we reasoned that if testosterone does not primarily cause dominant status-seeking, but instead, in non-threatening environments, promotes more agreeable and reputable status-enhancing behaviors [15, 16], participants in the testosterone (vs placebo) group should show a larger audience effect. Irrespective of whether testosterone would increase or decrease prosocial behavior under the audience effect, we also predicted that testosterone’s effects will be associated with changes in the efficiency of PE-based value updating (α in RL), choice consistency (tau in RL), and evidence necessary for making a decision (threshold parameter in DDM).

Previous research has suggested that testosterone possibly modulates social behavior through both androgenic and dopaminergic pathways and that testosterone effects on status-seeking behavior are moderated by CAG repeat polymorphism of the androgen receptor [27], and DAT1 polymorphism of the dopamine transporter [28]. We, therefore, examined whether these polymorphisms interact with testosterone administration effects on strategic prosociality. Furthermore, as research has shown that testosterone effects on status-seeking and decision-making are influenced by endogenous cortisol levels [29, 30], we examined whether testosterone administration effects interact with salivary cortisol levels measured at baseline and with cortisol reactivity to being watched. Moreover, as it has been suggested that sensitivity of dopaminergic pathways is heightened among highly dominant individuals [31] and that these individuals show more pronounced effects of testosterone administration [27, 32], we tested whether testosterone effects on strategic prosociality vary as a function of self-reported trait dominance [33]. Lastly, we sought to shed light on the motivational aspects [15, 34], of testosterone’s actions. We, therefore, conducted an exploratory post-hoc analysis that tested whether testosterone effects on strategic prosociality interact with self-reported value system, based on a questionnaire that captures the motivational bases of human attitudes and behavior [35, 36].

## Materials and Methods

### Participants

The study sample consisted of 192 healthy adult men aged between 18 and 40 years (*M* = 24.89, *SD* = 4.08). The sample size was determined based on previous testosterone administration studies [8, 13, 32] and our pilot study. In the applied linear mixed models, our sample size gave us 90% power to detect three-way and 86% power to detect 4-way interaction effects of size *f* ≥ 0.15. Using Monte Carlo simulation [37], we also calculated the probability of detecting a significant effect in the generalized linear mixed models (GzLMM) given our sample, experimental design, and the expected effect size based on previous research (*B* = 0.231) [16, 28]. The probability of a significant 3-way interaction was 93.68% (95% *CI* [89.08, 98.28]) and the probability of a significant 4-way interaction was 88.40% (95% *CI* [86.87, 89.31]. Participants were recruited via flyers placed around university campuses and online advertisements. The exclusion criteria comprised a history of neurological or psychiatric disorders, endocrine or other internal diseases, substance dependence, body mass index outside the healthy weight range (18.5 - 24.9), and the use of steroids. Only male participants were included as testosterone metabolism is subject to sex differences and the pharmacokinetics of topical administration of testosterone are unclear in women [38]. Two participants were excluded from the original sample *N=*192 because they continually clicked on the same response key irrespective of changing stimuli and reward probabilities for more than 80% of the block trials, and thus were classified as non-compliant. This led to a final sample size *N =* 190. All participants gave written consent and received a financial reward for their participation consisting of a flat fee and a bonus based on their task performance. All procedures were approved by the Ethics committee of the Medical University of Vienna and conducted following the latest revision of the Declaration of Helsinki [39]. No side-effects or adverse events were reported during or after the experimental sessions.

### Procedure and experimental design

Testing took place in groups of three to five participants, who were seated individually in small cubicles within the same testing room. All experimental sessions started between 01:00 and 02:30 p.m. First, a buccal smear sample for CAG repeat and DAT1 polymorphisms analysis was taken (see *SM*: *Supplementary information on the analysis of genetic data*). 20 min after arrival, participants were asked to drool 2 mL of saliva into a polyethylene collection tube. All salivary samples were frozen on-site and stored at - 30 °C until analysis. Afterward, participants were administered topical testosterone or placebo gel in a double-blind between-subjects design with random group allocation. Those allocated to the testosterone group received a single dose of testosterone gel, containing 150 mg testosterone [Androgel®]; participants in the placebo group received an equivalent amount of placebo gel. The only difference between the testosterone and placebo gel was that the placebo gel did not contain testosterone. Participants rubbed the gel onto their upper arms and shoulders using disposable latex gloves. Gel administration was followed by a 2-hour waiting period, during which participants remained in the laboratory premises, completed personality and demographic questionnaires, and were offered leisure-time reading materials. The testosterone dose and timing of the experiment were based on the previously established pharmacokinetic study of testosterone gel preparations in healthy young males [38]. One hour and 50 minutes after the gel application, participants provided a second saliva sample and subsequently began the experimental task (see Figure 1). Two more saliva samples were taken during the course of the study: 20 and 60 minutes after the end of the experimental task. After data collection was complete, saliva samples were analyzed by liquid chromatography-tandem mass spectrometry.

Participants performed the experimental task under one of the two randomly assigned between-subject conditions: either in private or when being observed. In the private condition, participants were informed that their performance is completely anonymous and no one (including the experimenter) would know how much money they would earn for themselves and the charitable organization. In the observed condition, two female observers, introduced as NGO association representatives entered the room and watched participants perform the task. The observers were seated at a desk with a laptop and had an equal view of all the participants. In addition, when participants were observed, a red frame was shown on their computer‘s screen and the displays of the participants’ computers were transmitted onto the observers’ laptop screens.

Participants were thus randomly assigned into four experimental groups corresponding to the levels of two between-subject factors: (1) treatment (testosterone/placebo) and (2) visibility (observed/private). These groups did not differ in age, trait dominance, basal hormone levels, or distribution of AR CAG and DAT1 genotype (see *SM: Table S1*).

### Prosocial learning task

Participants performed a probabilistic reinforcement learning task [25], where they could earn rewards either for themselves (self condition) or for an NGO of their choice (other condition). On each trial, participants were presented with two abstract symbols, one associated with a high (75%) and the other with a low (25%) reward probability. These contingencies were not instructed but had to be learned through trial and error. Participants selected a symbol by a button press and then received feedback on whether they obtained points or not. This way participants learned which symbol to choose to maximize the rewards in the long run. The points were converted to monetary rewards at the end of the experiment. Participants completed 6 blocks, 3 blocks in self and 3 blocks in the other condition. Each block started with a new pair of symbols and consisted of 16 trials/choices. In the self condition, the blocks started with “YOU” displayed and had the word “YOU” at the top of each screen. In the other condition, the blocks started with “NGO” displayed and had the word “NGO” at the top of each screen. The order of the blocks was pseudo-randomized so that the same recipient block did not occur twice in a row, and that half of the participants’ sample started the task with the *self* condition and the other half with the *other* condition. At the end of the experimental task, participants could choose the recipient of the money they earned in the other condition from a list of 6 different charities.

Immediately after completing the task, participants were asked to fill in a post-task questionnaire to estimate their subjective perception of being watched. The participants were asked the question: “Did you feel that you were being watched while performing the task?” The answers were classified into three categories: 1 (Not at all), 2 (Moderately), and 3 (Strongly).

### Statistical analysis of correct choices

Statistical analysis was performed using R statistical language [40]. We analyzed the treatment (testosterone/placebo) x visibility (observed/private) x recipient (self/other) interaction effect on correct choice using generalized linear mixed models (GzLMM) with binomial distribution and logit link function. The correct choice was defined as choosing the symbol with a higher reward probability. The participant’s identity was modeled as a random intercept effect and the within-subject factor recipient (self/other) was entered as a random slope.

To examine whether the effects of testosterone on correct choice varied as a function of trait dominance, CAG repeat, DAT1 polymorphism, baseline cortisol, cortisol reactivity, and personal value orientations, we added these variables separately as predictors in interaction with the other factors specified in the above GzLMM (See *SM* for information on the used R packages).

### Reinforcement learning drift-diffusion modeling

To uncover the cognitive computational processes underlying our learning task, we performed modeling analysis under the joint reinforcement learning drift diffusion model (RLDDM) framework [24, 41]. In essence, RLDDM bridges RL, which typically models choices, and DDM, which commonly models response times (RT). This approach has been proven to provide more granularity than using RL or DDM alone [24]. We tested candidate models with a single learning rate (Rescorla-Wagner models) as well as models with dual learning rates. Together, we tested 6 candidate RLDDM models, and the winning model is described below (see *SM: Supplementary information on computational modeling* for full model description, estimation, and comparison procedures). The RL part of the winning RLDDM model was implemented with a dual learning rates reinforcement learning model, where both the learning rate for a positive prediction error and the learning rate for a negative prediction error were employed to update values (i.e., *V*(A) and *V*(B) for two-choice options) [42] (Equation (1); see also *SM: Supplementary information on computational modeling).* The DDM part of the winning RLDDM model was implemented via a non-linear transformation of the accuracy-codded value differences computed from the RL counterpart, to construct the trial-by-trial drift rates [24] (Equation (2); see *also SM: Supplementary information on computational modeling*). The winning model contained 14 parameters: 7 separate parameters for each between-subject condition (i.e., placebo/testosterone, private/observed), and differential parameters for the within-subject condition (i.e., other/self; see *SM: Table S4* for the parameter list and description). In all models, we simultaneously modeled participants’ choice and RT, separately for each between-subject condition (i.e., placebo/testosterone; observed/private). Model estimations of all candidate models were performed with hierarchical Bayesian analysis (HBA). HBA was particularly useful when the number of trials was limited (here 16 trials per block) because in HBA, group-level and individual-level parameters were mutually informing each other during model estimation. All models reached convergence (see *SM: Supplementary information on computational modeling*).

### Statistical analysis of model parameters

The drug treatment (testosterone/placebo) x visibility (observed/private) x recipient (other/self) effect on the extracted free parameters was analyzed using GzLMMs analogous to the analysis of the correct choice. Due to the non-normal distribution of residuals, gamma distribution with a log link function was used for the parameter analyses. Finally, we tested whether the RLDDM parameter estimates, which were affected by the interaction of the drug treatment, visibility, and recipient could explain the differences observed in the behavioral prosociality measure. To do so, we conducted multiple linear regressions with the difference in the number of correct choices made for other and self (*prosociality index*) as a dependent variable and the differences in the RLDDM parameter estimates (αneg_other_-αneg_self_, τ_other –_ τ_self_, threshold_other_-threshold_self,_ drift-scaling_other_-drift-scaling_self_) as separate predictors. Bonferroni correction for multiple comparisons was used.

## Results

### Manipulation check

In the testosterone, compared to the placebo group, we observed higher salivary testosterone levels 110 minutes after gel administration, and this difference remained stable until the end of the experiment (drug treatment x time: *F*(3, 554.82) = 48.00, *p* = .001, *R^2^* =.737), see *Figure S1* and *SM: Supplementary analysis of hormone data* for analysis details). Analysis of the subjective ratings of being watched showed that participants in the observed condition felt watched to a greater extent than in the private condition (*Χ*^2^ (2, *N*=190) = 114.49, *p <*. 001), and that testosterone administration did not influence the perception of being watched (*Χ*^2^ (2, *N*=190) = 0.006, *p =*. 997). The degree to which the participants felt watched, was positively associated *r* (185) = .177, *p* = .016) with cortisol reactivity to the visibility manipulation (Δ cortisol levels 20 minutes after the end of the task - cortisol levels immediately before the task).

### Testosterone eliminates the audience effect

The three-way interaction of the factors drug treatment (P/T), visibility (private/observed), and type of recipient (self/other) predicted the number of correct choices (i.e., options that have higher reward probability) the participants made (*OR =* 0.94, *CI* = [0.89, 1.00], *p* = .043; Figure 2A). Follow-up analysis using treatment contrasts showed that participants in the placebo group showed more prosocial behavior, as indicated by relatively more correct prosocial choices when being watched compared to the private setting in which they were not watched (recipient x visibility interaction in the placebo group: *OR =* 1.43, *CI* = [1.01, 2.02], *p* = .042). Supporting our prediction based on the social dominance hypothesis, this audience effect was absent in the testosterone group (recipient x visibility interaction in the testosterone group: *OR* = 0.87, *CI* = [0.62, 1.22], *p* = .418). Specifically, when participants were observed, testosterone, compared to placebo, reduced the number of correct choices made for another (*OR =* 0.69, *CI* = [0.50, 0.94], *p* = .019, Figure 2C). The number of correct choices made for self, however, was not influenced by the drug treatment (*OR =* 0.98, *CI* = [0.75, 1.28], *p* = .875), visibility (*OR =* 1.00, *CI* = [0.78, 1.30], *p* = .982, or their interaction (*OR =* 0.88, *CI* = [0.61, 1.28], *p* = .509, Figure 2B).

**Figure 2.**
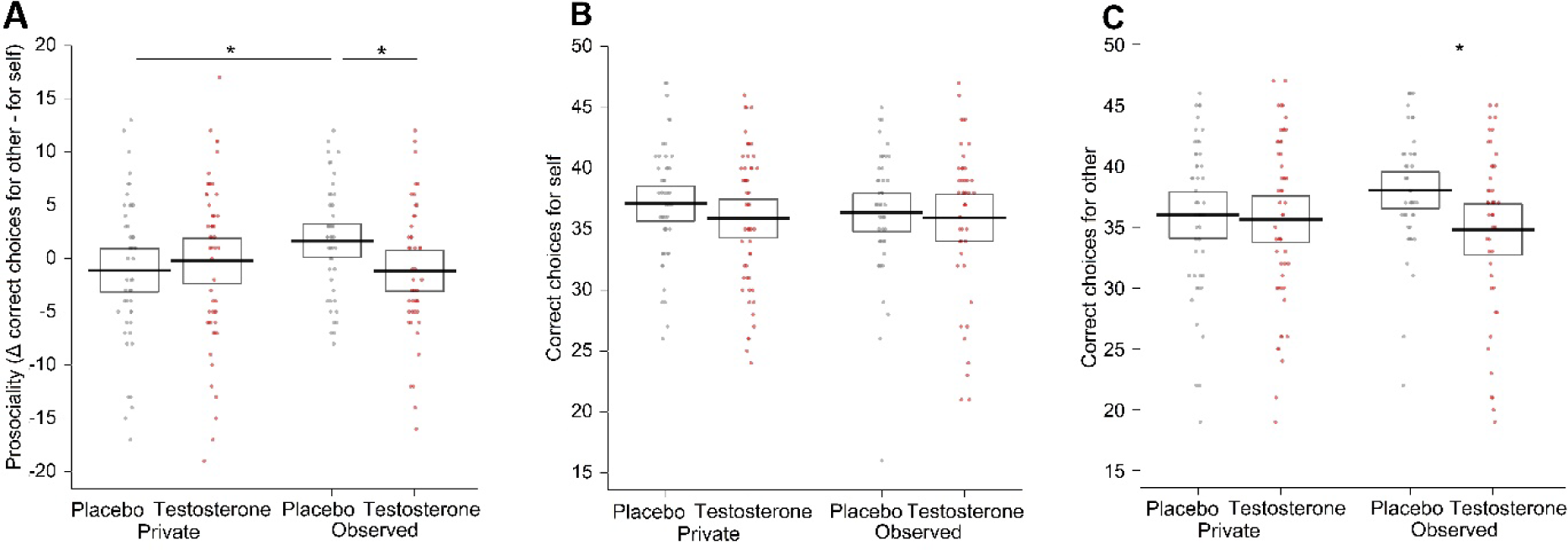
Differences in the number of correct choices. *(A)* Three-way interaction of the factors treatment x recipient x visibility. Participants in the placebo group behaved more prosocially (as captured by the prosociality index = correct choices for other – correct choices for self) when being observed than in privacy. Exogenous testosterone eliminated this audience effect. Note that the analyses were performed with raw data in the 2 x 2 x 2 factorial design; the plotted difference score (other minus self) is to improve readability and interpretability. Breaking down the three-way interaction by using treatment contrasts showed that there was no significant effect of the drug treatment and visibility factors, or their interaction, on the number of correct choices made for oneself *(B)*. On the other hand, testosterone, compared to placebo, decreased the number of correct choices made for the NGO when being observed *(C)*. Dots represent the data of individual participants, lines represent mean values per group, and boxes 95% confidence intervals.

### Behavior is best explained by a reinforcement learning drift diffusion model with dual learning rates

Next, we sought to uncover the computational mechanisms underlying the experiment-condition-specific behavioral differences on a trial-by-trial basis. The winning model (winning over five other candidate models; see *Materials and Methods*, *SM: Model selection and validation, and Table S3*) entailed combined RL and DDM components, and thus simultaneously predicted individuals’ choices and RTs (see *SM: Table S4* for a complete list of parameters and their description). The RL component section predicted participants’ learning behavior via the value updates through the computation of PEs with separate learning rates for positive and negative PEs (i.e., α^posPE^ and α^negPE^ Equation (1)). In other words, the model that best accounted for the data assumed a differential speed of learning with and without positive feedback:

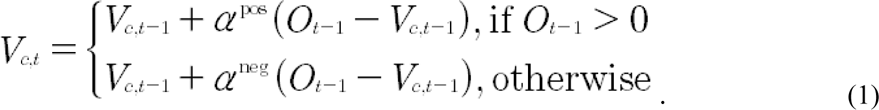

where *0*_*t*−1_ denotes the outcome, and *V*_*t*−1_ the subjective value of choice *c* at trial *t* − 1.

In addition, the DDM component predicted RTs by assuming an evidence accumulation process (as quantified by the drift rate; decisions were made when the evidence reached a certain threshold [41]). Importantly, the marriage between RL and DDM allowed a fine-grained investigation into how the drift rate (*v_t_*) was shaped by the value difference between two symbols at the trial-by-trial level (Equation (2); *S*, a non-linear transformation function; *v_scaling_*, a weight parameter that maps accuracy-coded value difference into the drift rate [24]).

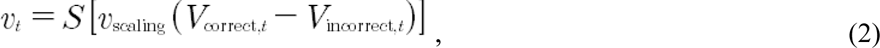

We fitted all candidate models (see *Materials and Methods* and *SM: Supplementary information on computational modeling*) under the hierarchical Bayesian estimation scheme [43] to incorporate both group-level commonality and individual differences, according to our task design (effects of drug treatment (P/T), visibility (private/observed), and type of recipient (self/other)).

### Testosterone’s impact on strategic prosocial behavior is associated with choice consistency

Next, we investigated which RLDDM parameters of our validated winning model are associated with the effects found in the behavioral analysis of the correct choice. As a first step, we tested the parameters for the 3-way interaction effect of drug treatment, visibility, and type of recipient. In the second step, we examined whether the parameters that showed a three-way interaction effect of our experimental manipulation predict behavioral prosociality (the difference between correct choices made for others and self). Out of the five parameters (learning rate for positive PE, learning rate for negative PE, choice consistency, threshold, drift-scaling parameter), only choice consistency showed the requisite three-way interaction of our experimental manipulations (*B =* 0.98, *CI* = [0.97, 0.98], *p* < .001), and at the same time significantly predicted behavioral prosociality (Bonferroni correction for multiple comparisons, *B* = 3.82, *CI* = [2.64, 5.01], *p* < .001; Figure 3D). Specifically, participants in the placebo group displayed relatively higher consistency in choices made for the other (vs. self) when being observed than in privacy (recipient x visibility interaction in the placebo group: *B =* 1.09, *CI* = [1.05, 1.14], *p* < .001). On the contrary, in the testosterone group, observation, compared to privacy, decreased the consistency of choices made for the other (vs. self) (recipient x visibility interaction in testosterone group: *B* = 0.90, *CI* = [0.84, 0.98], *p* < .001; Figure 3B). When participants were observed, testosterone, compared to placebo, diminished the relative consistency of prosocial choices (recipient x treatment interaction in observed condition: *OR* = 0.91, *CI* = [0.84, 0.99], *p* = .025; for analysis of all RLDDM parameters, see *SM: Analysis of the RLDDM parameters and their association with prosocial behavior*).

**Figure 3.**
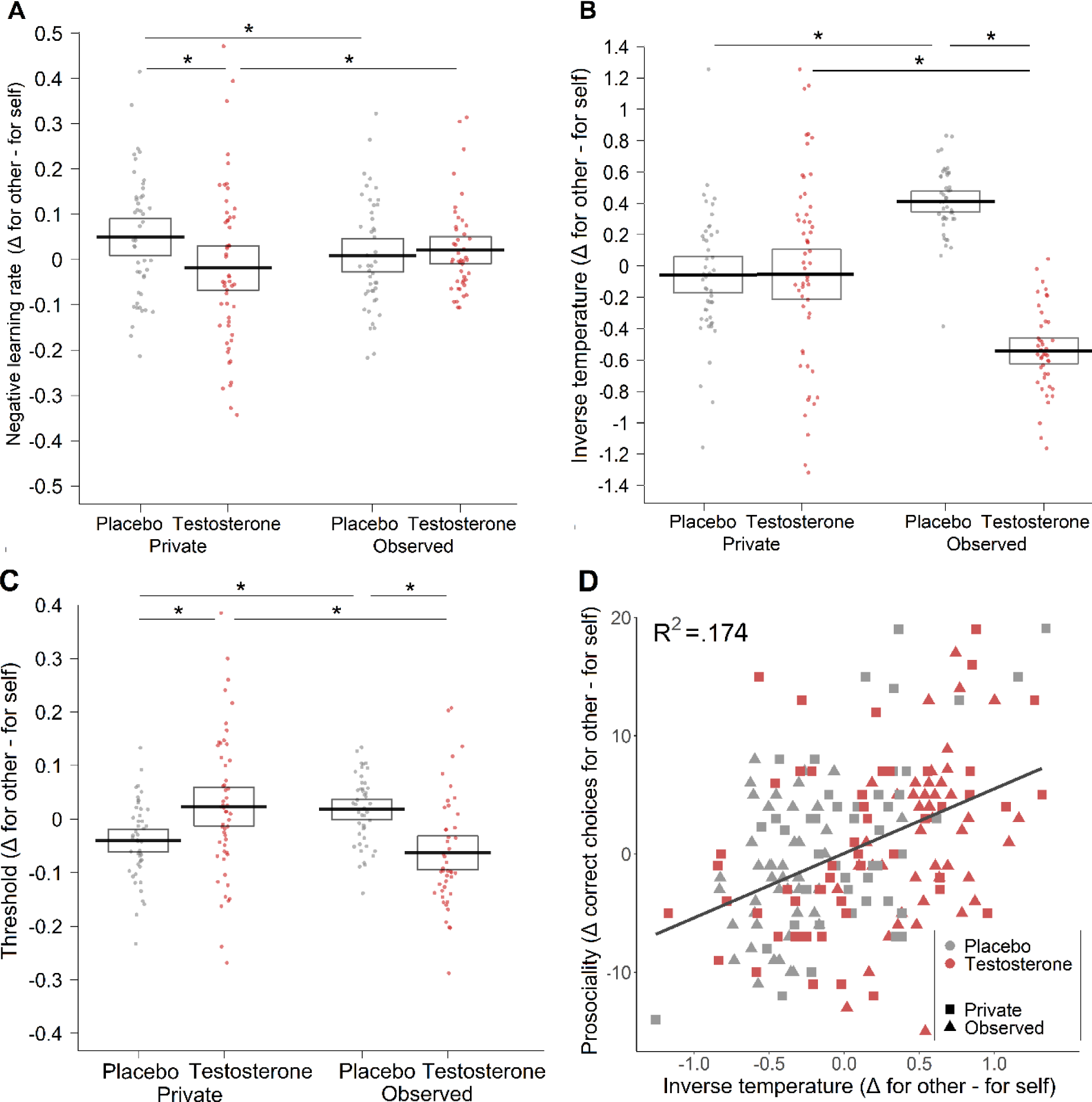
Differences in the parameters estimated by the reinforcement learning drift diffusion model (RLDDM). *(A)* In the placebo group, observation compared to privacy relatively decreased the prosocial learning rate for negative PE (i.e., the difference between α^negPE^ in the other condition and α^negPE^ in the self condition). Testosterone administration reversed the observation effect. The results suggest that for better performance in the task, a lower learning rate from negative PE is more suitable. *(B)* In the placebo group, observation compared to privacy, relatively increased the consistency of the prosocial choices. Testosterone administration reversed this audience effect. *(C)* In the placebo group, observation compared to privacy, relatively increased the DDM threshold for prosocial choices. Testosterone administration reversed the audience effect. *(D)* Inverse temperature parameter tau that captures choice consistency significantly predicted prosociality. Dots represent the data of individual participants, lines represent mean values per group, and boxes 95% confidence intervals.

Altogether, these results suggest that testosterone eliminates audience-dependent prosocial behavior by affecting choice consistency.

### Interaction of testosterone effects with trait dominance

In further support of the social dominance hypothesis, trait dominance interacted with testosterone’s effects on correct choice (recipient x drug treatment x visibility x trait dominance: *OR =* 1.04, *CI* = [1.01, 1.09], *p* = .026). In a follow-up analysis aimed at decoding this interaction, the continuous measure of dominance was replaced by a categorical variable with levels of high and low dominance, based on the median split of dominance scores (Med = 3.949). Decomposition of the four-way interaction revealed that testosterone reduced the number of correct choices made for others during observation specifically among men with high trait dominance (*OR =* 0.60, *CI* = [0.42, 0.87], *p* = .008) and this effect was weaker and non-significant among those with low dominance (*OR =* 0.77, *CI* = [0.55, 1.04], *p* = .132). Trait dominance did not significantly interact with the RLDDM parameters (all *p*s>.331 see *SM: Interaction of trait dominance with testosterone effects on RLDDM parameters)*.

### Interaction of testosterone effects with genetic polymorphisms and cortisol levels

CAG-repeat and DAT1 polymorphisms did not interact with the effects of testosterone on correct choice or RLDDM parameters (all *p*s>.091, see *SM: Supplementary information on the analysis of genetic data*). Furthermore, neither baseline cortisol levels, nor cortisol reactivity to visibility manipulation (Δ cortisol levels 20 minutes after the end of task-cortisol levels immediately before the task) interacted with the effects of testosterone on correct choice or RLDDM parameters (all *p*s >.052, see *SM: Supplementary analysis of hormone data, Figure S2, and Table S2 for details on cortisol levels*).

### Interaction of testosterone effects with personal value orientations

We next explored whether four principal value orientations (self-enhancement, self-transcendence, openness, conservation) measured using a self-report questionnaire [35,36] interacted with testosterone’s influence on the number of correct choices and choice consistency.

This revealed an interaction of *self-enhancement value orientation* with testosterone’s effect on the number of correct choices (recipient x visibility x administration x self-enhancement*: OR* = 0.65, *CI* = [0.48, 0.88], *p* = .006) and interaction of *conservation value orientation* and testosterone’s effects on choice consistency (recipient x visibility x administration x conservation: *B* = 0.60, *CI* = [0.17, 1.02], *p* = .006). The treatment contrast analysis of the interaction effect with self-enhancement showed that when testosterone participants were watched, a higher score in self-enhancement value was negatively associated with correct prosocial choices (*OR* = 1.29, *CI* = [0.21, 2.36], *p* = .008). In the placebo group, we did not observe such an association (*OR* = 1.11, *CI* = [0.89, 1.38], *p* = .349) and this difference between testosterone and placebo group was significant (*OR* = 0.72, *CI* = [0.55, 0.94], *p* = .018).

The treatment contrast analysis of the interaction effect with conservation value showed that when placebo participants were watched, the prosocial choice consistency was positively associated with conservation value (*B* = 1.29, *CI* = [0.22, 2.35], *p* = .019). However, after testosterone administration, the link between conservation value and prosocial choice consistency was abolished (*B* = −0.02, *CI* = [-0.94, 0.90, *p* = .969), but this difference between testosterone and placebo group did not reach significance (*OR* = 1.30, *CI* = [-0.10, 2.71], *p* = .069). No other interaction between value orientations and testosterone effect on choice behavior or RLDDM parameters was detected (all *p*s >.287 see *SM: Supplementary information on the questionnaire data)*.

### Learning parameters in relation to optimal learning rates

To gain a deeper understanding of how the learning parameters were related to the task performance in our experimental design, we performed a simulation study to identify optimal learning rates [44] (see *SM: Simulations of optimal learning rates*). In all conditions, both the posterior positive and negative learning rates were smaller with respect to the optimal ones (see Figure 4A, 4C). Crucially, to validate whether the choice accuracy corresponding to the posterior parameters in our winning model could capture key patterns in our behavior findings (i.e., posterior predictive check), we let our winning model generate synthetic data and analyzed the generated prosocial behavior (i.e., choice accuracy for other minus choice accuracy for self) in the same way as we analyzed the observed data. We found that results from the generated data (Figure 4B, 4D) greatly resembled the behavioral patterns reported in Figure 2A.

**Figure 4.**
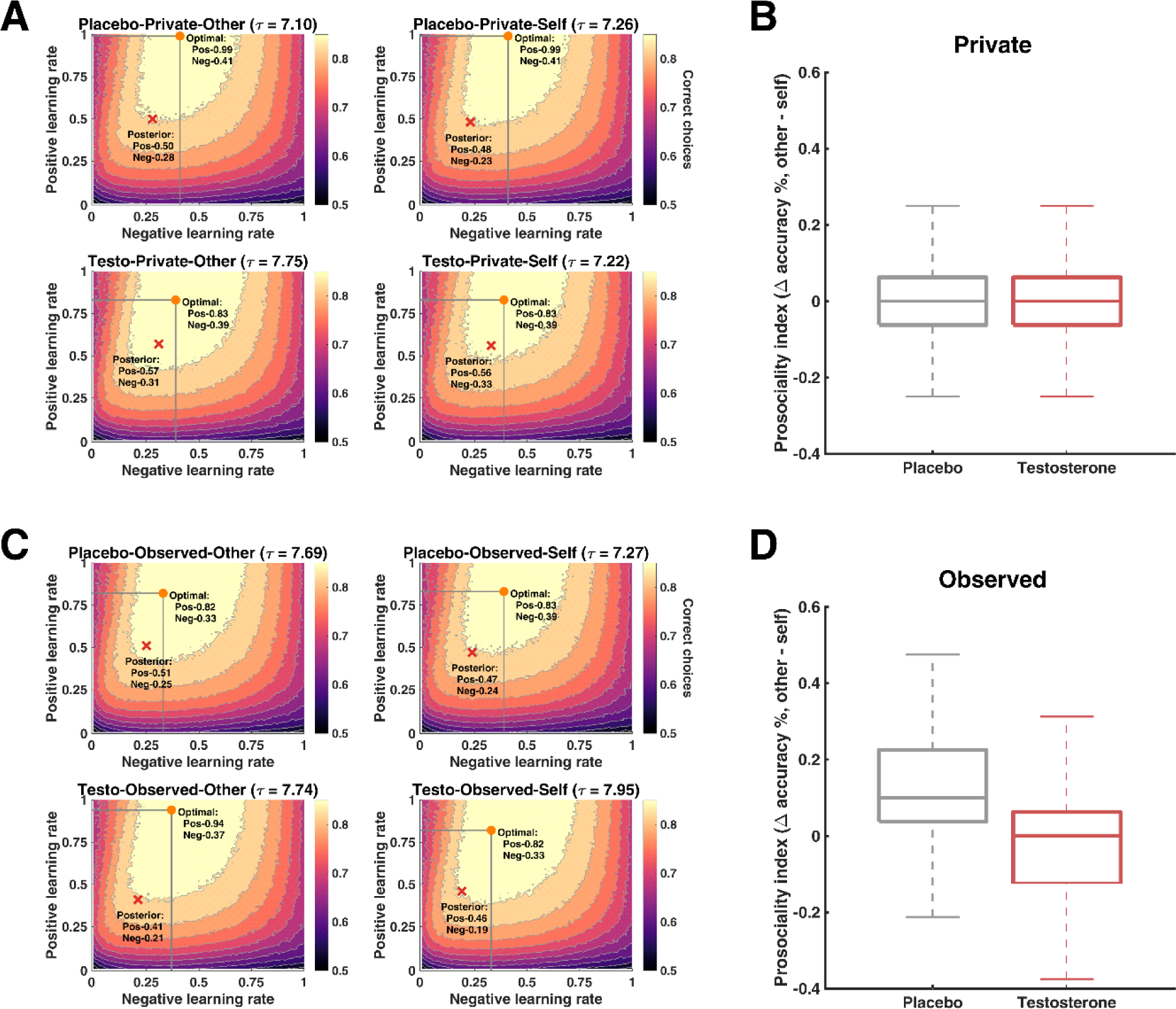
Optimal learning rates and posterior predictive checks. Posterior learning rates in relation to the optimal learning rates in the private *(A)* and observed *(C)* conditions. Orange dots represent the optimal combination between learning rates for positive and negative PE identified via simulation; red crosses indicate the posterior means of learning rates. The posterior learning rates were employed to perform posterior predictive checks for the main behavioral findings for the private *(B)* and observed *(D)* conditions. Simulated data from posteriors were analyzed in a similar fashion as the real data and the model prediction largely matched our main behavioral effect (cf. Figure 2A).

## Discussion

Using pharmacological manipulation and a novel computational model integrating reinforcement learning with the drift diffusion modeling framework (RLDDM), we tested and characterized testosterone’s role in audience-dependent prosocial behavior. The results show that testosterone diminishes the typical audience effect present in the placebo condition. Computational modeling pinpoints this effect to a reduction in the extent to which the performance of prosocial (vs. selfish) choices is consistent with learned reward values. Moreover, the effects are more pronounced in participants with higher trait dominance. Taken together, these findings are in line with the social dominance hypothesis and are thus consistent with the notion that testosterone decreases submission to audience expectations, rather than promoting the strategic display of socially pleasing behavior [45, 46].

A growing body of evidence suggests that testosterone exerts its behavioral effects through the modulation of reward-related processes [19, 20]. However, to our knowledge, no study investigated the computational mechanisms underlying such effects. Using joint RLDDMs, we found that in the placebo group, observation (vs privacy) increased the relative consistency of prosocial choices. Testosterone administration eliminated this audience effect, making the performance of prosocial (vs. self-benefitting) choices less consistent with value computations. Low choice consistency means that individuals select options with non-maximal expected values, which is often referred to as exploratory behavior [47]. In environments with static reward probabilities, participants can maximize their reward by initially exploring which option tends to be more fruitful. Once learners discover the better option, exploration yields no benefit. One possible explanation of the present effect could therefore be that testosterone impaired individuals’ ability to adapt and control the amount of exploration. However, our data do not indicate that testosterone affects exploration in general, as we did not find any testosterone influence on choice consistency in the private setting. Instead, the effects of testosterone appeared only in a situation where social status was at stake. Interestingly, research in monkeys has shown that socially challenging environments (vs socially isolated environments) increased the availability of D_2_ receptors [48], which play a role in choice consistency [49]. This effect was present in particular in dominant monkeys and absent in the subordinate ones. Such findings are in line with our results showing enhanced testosterone effects in individuals with high trait dominance. We, therefore, suggest that future studies should examine the potential mediating role of D2 receptors in testosterone’s effect on reward learning in social situations.

Alternatively, it could be speculated that the elimination of the audience effect by testosterone stems from the hormone’s ability to reduce fear in social situations. Indeed, earlier research shows that exogenous testosterone diminishes the physiological stress response to the presence of an observer [13] and has anxiolytic-like properties in humans and across species [10, 50]. However, we did not observe any interaction of the testosterone’s effect with cortisol levels measured at baseline or with cortisol reactivity, thus our data do not provide support for such an interpretation. To examine this topic further, studies on testosterone and audience effect should include more explicit measures of subjectively perceived stress and anxiety.

The present data also shows that testosterone administration substantially alters the relation between the audience effect and self-reported value orientations. *Self-enhancement* value, as measured here [35], is characterized by power, achievement, and the pursuit of one’s own interests, success, and dominance over others. We found that a higher emphasis on self-enhancement was related to a lower number of correct prosocial choices while being watched in the testosterone, but not in the placebo group. This finding extends the evidence that the hormone testosterone interacts with dominant personality traits. In contrast to self-enhancement, conservation value reflects a personal emphasis on conformity, security, tradition, and self-restriction [35]. We showed that in the placebo group, the degree to which one identifies with conservation value is positively associated with the consistency of prosocial choices made while being watched. Testosterone administration abolished this link between conservation value and prosocial choice consistency. In consideration of the conformity aspect of the conservation value, it is of interest that in Western societies, nonconforming people are perceived as having higher status and competence than those who conform to the social expectations [51]. Thus, these findings are suggestive of the interpretation that by inducing a non-confirming attitude, testosterone reaches its principle goal - to be observed as having high status and competence^17,46^. We note though that the analyses of value orientations are exploratory and were performed post-hoc, and thus need independent confirmation.

Variability in dominance, conservation, and cultural differences in social status attaintment, can also account for the results of another recent study, which was conducted among Chinese students and showed that testosterone enhanced audience effects [16]. Indeed, contrary to Western society, in Eastern cultures, high social status is associated with increased self-restriction and other-orientation [52]. Consistently, individuals from Western and Eastern cultures differ in their self-construal, i.e., in the way they define the self in relation to others, with, for example, European individuals construing themselves as being more independent, or less interdependent, than Asian individuals [53]. Interestingly, research has shown that acute testosterone changes in men are positively associated with aggressive behavior for those with more independent self-construals, whereas basal testosterone is negatively associated with aggression when individuals have more interdependent self-construals [54]. Moreover, the cultural differences in independent vs. interdependent social orientations have been linked to polymorphisms in the dopamine D4 receptor gene [53], implying a putative biological mechanism that could explain cultural differences in testosterone effects. Finally, contrasting results may also stem from differences in the applied methods. While Wu et al. [16] used a modified dictator game, where participants were explicitly asked whether they want to donate a certain monetary amount to charity, in the present task, the donation to charity was determined indirectly by the participant’s performance in a reinforcement learning task. Our paradigm thus presents a more implicit measure of prosociality. For these reasons, we suggest future studies of how testosterone affects status-seeking behavior should focus on exploring these cultural differences, specifically by including measures of value orientations, self-construal, implicit behavioral tasks, as well as assessments of dopamine receptor polymorphisms.

Our results are, furthermore, in line with studies showing that testosterone decreases deception [55-57]. Further research is, however, needed to determine whether testosterone reduces lying per se, or only in situations where dishonest behavior may be considered “cheap”, dishonorable, and lower the subject’s feelings of pride and self-image [55].

There are also some limitations inherent to the methodology of our study. Due to the sex differences in testosterone metabolism and unknown pharmacokinetics following the topical administration of testosterone in women [38], the study included only male participants. Hence, the generalization of these findings to females requires further investigation. Furthermore, the observers in our study were exclusively female. Future research will also be required to more systematically test whether testosterone’s influence on the prosocial audience effect is sensitive to the gender, number, and salience of the present observers. Nevertheless, our survey data suggest that the salience of the observers used in our study was representative of a wide range of social contacts and that the subjective feeling of being watched scaled with an established stress biomarker.

In conclusion, we conducted a multifaceted examination of the computational, endocrinological, and genetic mechanisms underlying audience effect and showed that testosterone reduced strategic prosocial learning through impairment of choice consistency. These findings provide evidence that in the Western student sample, testosterone abolishes audience effects, and therefore does not foster the seeking of social leadership by reputational politics. The present study is the first to specify testosterone’s role in reward processing by revealing that testosterone impacts status-seeking through modulation of how the learned reward values are expressed in behavior.

## Acknowledgments

We thank Felix Dörflinger and Maximilian Kathofer for their dedicated assistance with data collection. We also thank Shawn Geniole for his helpful feedback.

## Author Contributions

H.H.K. and L.Z. share the first authorship of this article. H.H.K, C.E., and C.L. designed research;

H.H.K. performed research; H.H.K. and L.Z. analyzed data; H.H.K., L.Z., C.L., and J.v.H. wrote the paper.

## Supplementary Materials for

Other supplementary materials for this manuscript include the following: Data and codes can be accessed at: https://osf.io/qr4ve/

### Supplementary information on the use of statistical software and functions

Statistical analysis was performed using the R statistical language [1] and the packages: lme4 [2] for construction of general linear mixed models (GLMM) and generalized linear mixed model (GzLMM); car package [3] for construction of general linear models and computation of *p-*values based on Type III Wald chi-square tests; sjPlot package [4] for post-hoc tests of significant three-way interactions including odds ratios (ORs) and 95% confidence intervals (95%CIs); yarrr [5] and ggplot2 [6] packages for construction of plots. Computational modeling was performed using Markov chain Monte Carlo with the statistical computing language Stan [7] while following the hBayesDM package [8]. Model comparison and evaluation were performed using the LOO package [9].

### Supplementary analysis of hormone data

#### The effect of drug treatment on hormone levels

After data collection was complete, saliva samples were shipped on dry ice to Dresden LabService GmbH led by Clemens Kirschbaum, Germany. Liquid chromatography-tandem mass spectrometry was used to determine the hormonal levels. To examine the change of the hormonal levels throughout the experimental session, testosterone, cortisol, and estradiol levels were analyzed using GLMMs with the fixed factors drug treatment (testosterone/placebo), visibility (observed/private), time (baseline/1 h 50 min after drug treatment/20 min after the end of the task/60 min after the end of the task), and participant’s identity as a random intercept. Due to the non-normal distribution of residuals, hormonal data were log-transformed. Baseline hormonal levels did not significantly differ across experimental groups (all *p*s > .336, see Table S1). As expected, 1h 50 min after gel administration, we observed higher testosterone levels in the testosterone group (M_Sample2_ = 5014.10 pg/mL, 95%CI [3866.10, 6502.88]) compared to the placebo group (M_Sample2_ = 134.30, 95%CI [103.54, 175.92]; drug treatment x time: *F*(3,554.82) = 48.00, *p* < .001, *R^2^* =.737), a difference that remained stable until the end of the experiment (see Figure S1).

There were no effects of drug treatment on cortisol (drug treatment x time: *F*(3,520.6) = 1.246, *p* = .292) or estradiol levels (drug treatment x time: *F*(3,550.79) = 1.497, *p* = .214).

The observation condition did not significantly influence any hormonal levels (testosterone: visibility x time: *F*(3,554.82) = 2.447, *p* = .063; cortisol: visibility x time: *F*(3,520.6) = 0.900, *p* = 441; estradiol: visibility x time: *F*(3,550.79) = 0.796, *p* = .496).

#### Contamination of salivary samples

Out of a total of 192 baseline samples, we noted that 34 contained above-normal testosterone levels, atypical for normal young men (> 2000 pg/mL). All other baseline values were hormonally typical. The samples with abnormally high testosterone values appeared only in the participants, who later received testosterone treatment, not in the placebo group. Previous research [10, 11] described similar abnormally high testosterone levels and attributed it to the testosterone contamination of the common surfaces (e.g., doorknobs, keyboards), excluding the option of physiological contamination. Based on their recommendation, we implemented a cleaning protocol that included the wearing of disposable sterile gloves, and cleaning of keyboards, computer mice, tables, and doorknobs with an alcohol-based solution after each session. Although these precautions successfully prevented between-session contamination, we suspect that they still did not reliably impede within-session contamination of the saliva containers. For future studies, we, therefore, recommend even stricter sanitizing protocols and more careful handling of the saliva collection tubes and boxes, before, during, and also after sample collection.

In our sample, the abnormally high values were present only in the sessions where testosterone was administered, and this notwithstanding, the testosterone group showed a reliable testosterone increase after the drug administration in comparison to the placebo group. We, therefore, decided to retain the participants with contaminated baseline samples for the behavioral analyses, except for the analysis that includes baseline testosterone levels.

#### The effect of salivary testosterone levels on the correct choice

To examine whether salivary testosterone levels measured in the saliva samples taken before the start of the experimental task (i.e., 2 hours after the drug administration) predicted participants’ behavior, we included the mean-centered log-transformed testosterone levels as a predictor in interaction with the factors recipient and visibility to the GzLMM of the correct choice. The analysis did not reveal a main effect of testosterone levels (*OR =* 0.99, *CI* = [0.96, 1.03], *p* = .624) or a significant interaction effect on correct choice (recipient x drug treatment x testosterone levels: *OR =* 1.02, *CI* = [0.99, 1.05], *p* = .162). The absence of a significant association between salivary testosterone concentrations and behavior is in line with studies that point out that although salivary testosterone measurements are correlated with the hormone concentration in serum, they do not precisely track the availability of free serum testosterone after transdermal application [11, 12]. Thus, although the between-groups comparison of saliva testosterone provides a manipulation check of topical drug administration, for the analysis of the relationship between behavior and post-administration hormonal levels on the individual level, salivary testosterone measures may presently lack sensitivity.

#### Interaction of salivary cortisol levels with testosterone effects on the correct choice and RLDDM parameters

To examine whether cortisol levels interacted with testosterone’s effect on correct choice and reinforcement learning drift diffusion model (RLDDM) parameters, we separately added baseline cortisol levels and cortisol reactivity as predictors to our main analysis. The cortisol reactivity was defined as a difference between cortisol levels detected in the saliva sample taken 20 minutes after the end of visibility manipulation and cortisol levels from the sample taken immediately before the start of the paradigm. The values were mean-centered and entered as a predictor in interaction with the other factors (recipient, drug treatment, visibility) to the GzLMM of correct choice and the GzLMMs of the RLDDM parameters with log link function (see *Materials and Methods* in the main manuscript text). The analysis revealed no significant interaction of baseline cortisol levels with testosterone effect on correct choice (recipient x drug treatment x visibility baseline cortisol: *OR =* 1.02, *CI* = [0.97, 1.07], *p* = .385), α^posPE^ (recipient x drug treatment x visibility baseline cortisol: *B =* 1.01, *CI* = [0.98, 1.04], *p* = .549), α^negPE^ (recipient x drug treatment x visibility baseline cortisol: *B =* 0.97, *CI* = [0.94, 1.00], *p* = .052), choice consistency (recipient x drug treatment x visibility baseline cortisol: *B =* 1.00, *CI* = [0.99, 1.01], *p* = .583) or decision threshold (recipient x drug treatment x visibility baseline cortisol: *OR =* 1.00, *CI* = [1.00, 1.0], *p* = .902).

Similarly, the analysis revealed no significant interaction of cortisol reactivity with testosterone effect on correct choice (recipient x drug treatment x visibility x post-task cortisol: *OR =* 1.04, *CI* = [0.99, 1.09], *p* = .133), α^posPE^(recipient x drug treatment x visibility x post-task cortisol: *B =* 1.01, *CI* = [0.99, 1.03], *p* = .454), α^negPE^ (recipient x drug treatment x visibility x post-task cortisol: *B =* 1.01, *CI* = [0.99, 1.01], *p* = .078), choice consistency (recipient x drug treatment x visibility x post-task cortisol: *B =* 1.00, *CI* = [1.00, 1.01], *p* = .249) or decision threshold (recipient x drug treatment x visibility x post-task cortisol: *B =* 1.00, *CI* = [1.00, 1.00], *p* = .996).

### Supplementary information on computational modeling

#### Rescorla-Wagner (RW) model

We started with the simple Rescorla-Wagner [13] model as our baseline model. On each trial, the value (*V*_c,t_) of the chosen option was updated with the reward prediction error (RPE):

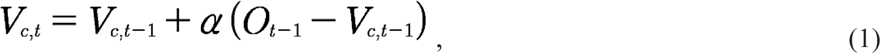

where *O*_t-1_ was the received outcome, and α (0 < α < 1) denoted the learning rate.

#### Dual learning rates (DLR) model

Previous studies have reported differences in learning following positive and negative PEs [14, 15] and these differences have been linked to the distinctive roles of striatal D1 and D2 dopamine receptors in segregated cortico-striatal pathways [16]. Furthermore, analyses of genetic polymorphisms of DARPP-32 that predict choice behavior associated with a positive outcome, and the DRD2 gene predicting avoidance of choices associated with negative outcomes, support the notion that independent dopaminergic mechanisms contribute to learning from positive and negative feedback [17]. RL models with dual learning rates have been successful in capturing this asymmetric learning effect [18,19], including studies in social neuroscience examining prosocial behavior [20]. Hence, we tested a dual learning rates model on top of the RW model:

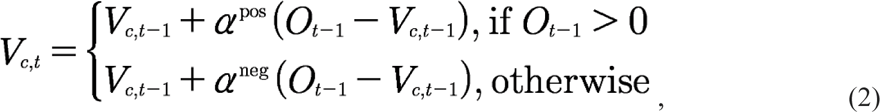

where α^posPE^ and α^negPE^ were the learning rates for positive and negative RPEs, respectively.

In both RW and DLR models, action values were converted to action probabilities using the softmax function. Let A and B be the choice symbols per trial, the probability of choosing A was computed via the difference between *V*(A) and *V*(B):

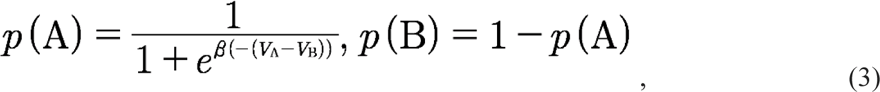

where β (β > 0) was the inverse temperature that represented choice consistency. Higher β indicated that individuals’ choices were more consistent with their value computation, where lower β indicated that individuals behaved more randomly. The action probability was then used to model participants’ choice data with a categorical distribution:

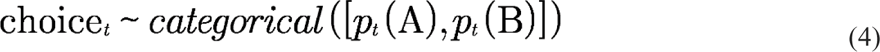

It is worth noting that our winning model with differential learning rates for positive and negative PEs is in line with previous studies on learning and decision-making that report asymmetric learning effect [14, 15, 18, 19]. The theoretical interpretation of these differences, however, is mixed in the literature. While some studies reported enhanced learning after positive PE compared to negative and related this feature to optimism bias [21], others report a higher learning rate for negative than for positive PEs, interpreted as possibly reflecting risk aversion [14]. Our findings in this regard may contribute to this debate such that learning update is much quicker after receiving positive feedback, especially when the feedback is associated with appetitive stimuli (e.g., monetary reward). This is likely to be generalized to situations with relatively positive feedback, rather than the actual positive feedback *per se*. For instance, in aversive learning, receiving no feedback (e.g., a neutral outcome) is, on a relative scale, more positive than actual negative feedback (e.g., an electric shock). Learning rates for such no-feedback events have been shown to be higher than those for stimuli with negative feedback, including learning under social contexts [20].

#### Drift diffusion model (DDM)

The drift diffusion model (a.k.a., diffusion decision model [22]) was a widely used computational framework to model individuals’ response times (RTs). In its canonical expression, DDM contained four parameters, namely, the drift rate (v; v > 0), the initial bias (z; z > 0), non-decision time (T; 0 < T min (RT)), as well as the decision threshold (a). For simplicity in learning tasks with abstract symbols, the initial bias z was fixed at 0.5. Trial-by-trial RTs were distributed according to the Wiener first passage time (WFPT [23]):

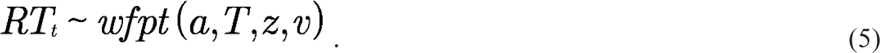

#### Reinforcement learning drift diffusion model (RLDDM)

In value-based decision-making, individuals’ RTs may vary as the function of trial-by-trial valuation, such that the larger the value difference between choice alternatives, the faster the RT. Therefore, a joint reinforcement learning drift diffusion model (RLDDM) framework has been proposed [15,24], bridging RL and DDM. This approach provides more granularity than using RL or DDM alone [24]. In essence, the drift rate in DDM was characterized by the accuracy-coded value differences computed from the RL counterpart. This way, the drift rate was no more a constant parameter throughout the entire experiment, instead, it varied across trials (i.e., *v_t_*, instead of *v*) according to the values computed from RL updates (in the present study, RW or RP). In the simplest RLDDM, trial-by-trial drift rates were constructed via a linear function of value difference:

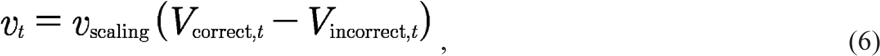

where *v*_scaling_ (v_scaling_ >0) was the scaling parameter that quantified the impact of value difference. Note that we employed stimulus coding in our RLDDM, so that in Equation (6), the drift rate was always a function of the value difference between the correct (i.e., more rewarding, 75% reward probability) and the incorrect options (i.e., less rewarding, 25% reward probability), rather than between the chosen and unchosen options.

#### Reinforcement learning drift diffusion model with non-linear transformation (RLDDM-nonlin)

There is evidence that a non-linear mapping between value difference and the drift rate could better capture individuals’ RTs as opposed to a linear transformation [24]. This is likely because non-linear functions may provide more sensitivity, akin to the softmax function in choice models. We thus implemented an RLDDM-nonlin following:

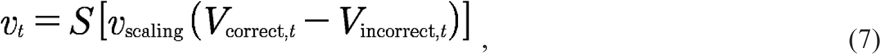

with

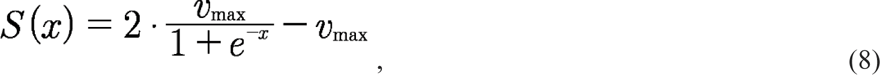

where *S*(x) was a non-linear sigmoid function centered at 0, that convert x to lie between –*v*_max_ and *v*_max_ (*v*_max_ > 0). It is worth noting that *v*_max_ only affected the maximum value of the drift rate, whereas *v*_scaling_, as in Equation 6, established the trial-by-trial mapping between the value difference and the drift rate.

In both RLDDM and RLDDM-nonlin, all other DDM parameters (i.e., *a*, *T*, *z*) were identical to the canonical DDM model, and RTs were distributed with *wfpt* using trial-by-trial drift rate (*v_t_*):

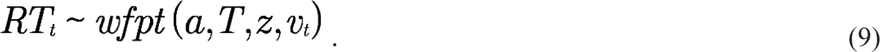

Note that, in all candidate models (Table S3), we introduced differential parameters for the within-subject condition of our experiment, namely, all parameters were separately modeled for the “self” and the “other” conditions.

#### Model estimation

The model estimation and model selection procedures were largely similar to [25]. Hence, below we echoed these procedures from [25] to enhance reproducibility, with modifications that were specific to the current study.

In all models, we simultaneously modeled participants’ choice and RT, separately for each between-subject condition (i.e., placebo vs. testosterone; observed vs. private). Model estimations of all candidate models were performed with hierarchical Bayesian analysis (HBA) [26] using the statistical computing language Stan [7] in R. Stan utilizes a Hamiltonian Monte Carlo (HMC; an efficient Markov Chain Monte Carlo, MCMC) sampling scheme to perform full Bayesian inference and obtain the actual posterior distribution. We performed HBA rather than maximum likelihood estimation (MLE) because HBA provides much more stable and accurate estimates than MLE [24]. Following the approach in the “hBayesDM” package [8] for using Stan in the field of reinforcement learning, we assumed, for instance, that a generic individual-level parameter *ϕ* was drawn from a group-level normal distribution, namely, *ϕ* ∼ Normal (*μ_ϕ_*, *σ_ϕ_*), with *μ_ϕ_* and *σ_ϕ_*. being the group-level mean and standard deviation, respectively. Both these group-level parameters were specified with weakly-informative priors [26]: *μ_ϕ_* ∼ Normal (0, 1) and *σ_ϕ_*.∼ half-Cauchy (0, 1). This was to ensure that the MCMC sampler traveled over a sufficiently wide range to sample the entire parameter space.

Appropriate parameter transformations were applied to double-bounded parameters (e.g., learning rate, [0, 1]) with the inverse probit function (i.e., the cumulative distribution function of the standard normal distribution), and single-bounded parameters (e.g., drift rate, (0, +∞)) with the soft-plus function (i.e., ln(1 + e^x)), respectively.

In HBA, all group-level parameters and individual-level parameters were simultaneously estimated through the Bayes’ rule by incorporating behavioral data. We fit each candidate model with four independent MCMC chains using 1,000 iterations after 1,000 iterations for the initial algorithm warmup per chain, which resulted in 4,000 valid posterior samples. The convergence of MCMC chains was assessed both visually (from the trace plot) and through the Gelman-Rubin ^R^^ Statistics [27]. ^R^^ values of all parameters were smaller than 1.05 in the current study), which indicated adequate convergence.

#### Model selection and validation

For model comparison and model selection, we computed the Leave-One-Out information criterion (LOOIC) score per candidate model [28]. The LOOIC score provides the point-wise estimate (using the entire posterior distribution) of out-of-sample predictive accuracy in a fully Bayesian way, which is more reliable compared to information criteria using point-estimate (e.g., Akaike information criterion, AIC; deviance information criterion, DIC). By convention, a lower LOOIC score indicates better out-of-sample prediction accuracy of the candidate model. We selected the model with the lowest LOOIC as the winning model. We additionally performed Bayesian model averaging (BMA) with Bayesian bootstrap [29]to compute the probability of each candidate model being the best model. Conventionally, the BMA probability of 0.8 (or higher) is a decisive indication.

Moreover, given that model comparison provided merely relative performance among candidate models [28], we then tested how well our winning model’s posterior prediction was able to replicate the key features of the observed data (a.k.a., posterior predictive checks, PPCs). Since we only found an effect in choice data, we performed PPCs only for choices (excluding RTs). To this end, we applied a one-step-ahead PPC [25, 30] that factored in participants’ actual action and outcome sequences to generate predictions with the entire posterior MCMC samples. Specifically, we let the winning model generate choices as many times as the number of MCMC samples (i.e., 4,000 times) per trial per participant, and we analyzed the generated data the same way as we did for the observed data. We then assessed whether these analyses could reproduce the behavioral pattern in our behavioral analyses (Figure 4B, 4D in the main text).

#### Simulations of optimal learning rates

To better understand and interpret the magnitude of the posterior learning rates, we performed simulations with grid approximation to obtain “optimal learning rates”, and then compared the estimated posterior learning rates in relation to these optimal parameters (Figure 4A, 4C in the main text). Because there were two learning rates (α^posPE^ and α^negPE^), to reduce complexity, we fixed the inverse temperature parameter to be the corresponding group-level posterior mean in each condition. For each simulation, we took a small grid per parameter (0:0.01:1) and computed the choice accuracy across 16 trials (identical to the main experiment) for each combination of the parameters. Each simulation was repeated 1000 times to obtain stable results. We then considered the parameters that gave the highest choice accuracy as the optimal learning rates.

#### Analysis of the RLDDM parameters and their association with prosocial behavior

Next, we examined whether the behavioral pattern found in the analysis of the correct choice would be associated with differences in the individual model parameters.

As a first step, we tested the parameters of our validated winning model for the 3-way interaction effect of drug treatment, visibility, and type of recipient. There was no significant 3-way interaction in the learning rate for positive PE (*B =* 1.03, *CI* = [1.00, 1.06], *p* = .063). The analysis of the learning rate for negative PE revealed a three-way interaction of drug treatment, visibility, and type of recipient (*B =* 0.94, *CI* = [0.90, 0.98], *p* = .003) so that the participants in the placebo group had a relatively lower negative learning rate for prosocial choices when being watched than in privacy (recipient x visibility interaction in the placebo group: *B =* 0.68, *CI* = [0.40, 0.96], *p* = .023). Conversely, in the testosterone group, observation (vs privacy) relatively increased the negative learning rate for prosocial choices (recipient x visibility interaction in testosterone group: *B* = 1.21, *CI* = [1.00, 1.40], *p* = .048; Figure 2A). Moreover, the analysis of the choice consistency (inverse temperature parameter tau, described also in the main text) likewise showed a three-way interaction (*B =* 0.98, *CI* = [0.97, 0.98], *p* < .001). Placebo group participants had relatively higher consistency in choices made for the other (vs. self) when being observed than in privacy (recipient x visibility interaction in the placebo group: *B =* 1.09, *CI* = [1.05, 1.14], *p* < .001). On the contrary, in the testosterone group, observation, compared to privacy, decreased the consistency of choices made for the other (vs. self) (recipient x visibility interaction in testosterone group: *B* = 0.90, *CI* = [0.84, 0.98], *p* < .001). When participants were observed, testosterone, compared to placebo, diminished the relative consistency of prosocial choices (recipient x treatment interaction in observed condition: *OR* = 0.91, *CI* = [0.84, 0.99], p =.025). In the private condition, there was no evidence for such an effect (recipient x treatment interaction in private condition: *OR* = 0.99, *CI* = [0.97, 1.02], p = .605). The analysis of the DDM threshold parameter revealed a three-way interaction as well (*B=* 1.01, *CI* = [1.00, 1.02], *p* < .001; Figure 2C). Placebo group participants had a relatively higher threshold for choices made for another (vs. self) when being observed than in privacy (recipient x visibility interaction in the placebo group: *B =* 1.03, *CI* = [1.01, 1.05], *p* < .001). Conversely, in the testosterone group, observation, compared to privacy, decreased the amount of information required for choices made for another (vs. self) (recipient x visibility interaction in testosterone group: *B* = 0.95, *CI* = [0.93, 0.97], *p* < .001). When participants were observed, testosterone, compared to placebo, decreased the relative threshold of prosocial choices (recipient x treatment interaction in observed condition: *B* = 0.95, *CI* = [0.93, 0.97], p < . 001). The analysis of the DDM drift-scaling parameter revealed a three-way interaction (*B* = 1.01, *CI* = [1.00, 1.02], p < .001). Participants in both placebo (recipient x visibility interaction in placebo group *B* = 0.93, *CI* = [0.91, 0.95], *p* < .001) and testosterone group (recipient x visibility interaction in placebo group *B* = 0.73, *CI* = [0.51, 0.95], *p* < .001) showed relatively lower drift scaling for choices made for another (vs. self) when being observed than in privacy. When participants were observed, testosterone, compared to placebo, decreased the relative drift scaling of prosocial choices (recipient x treatment interaction in the observed group: *B* = 0.91, *CI* = [0.90, 0.92], p < .001).

As a second step, we examined whether the RLDDM parameters that were impacted by testosterone administration predict behavioral prosociality, measured by the difference between correct choices made for other and self across the whole sample. Out of the five parameters, choice consistency (*B* = 3.82, *CI* = [2.64, 5.01], *p* < .001), and DDM threshold (*B* = 10.67, *CI* = [1.57, 19.76], *p* = .022) predicted prosociality, however, only choice consistency survived the Bonferroni correction for multiple comparisons (*p* <.01). Altogether, as reported in the main text, these results suggest that testosterone’s impact on strategic prosocial behavior (i.e., audience effect) is strongly linked to testosterone’s effect on choice consistency (inverse temperature parameter tau).

#### Analysis of the drift-scaling parameter and response times

As specified in equation (7), on each trial *t*, the drift rate *v_t_* was defined with a drift-scaling parameter, *v*_scaling_ that scales the value difference between the correct and incorrect symbol. Drift-scaling parameter affects the curvature of the function: smaller values lead to a more linear mapping between the value difference and the drift rate, and therefore less sensitivity to value differences.

Drift scaling is conceptually linked to the speed of integration and response times [22], we, therefore, tested whether drift-scaling parameter predicted response times and found a significant association (*B* = 0.97, *CI* = [0.95, 0.98], *p* < .001). However, contrary to correct choices, response times did not differ across experimental groups (drug treatment: *B* = 1.01, *CI* = [0.97, 1.05], p = .737; visibility: *B* = 0.98, *CI* = [0.94, 1.02], *p* < .265; recipient: *B* = 0.99, *CI* = [0.99, 1.00], *p* < .106; drug treatment x visibility x recipient: *B* = 1.00, *CI* = [0.99, 1.01], *p* < .806).

### Supplementary information on the analysis of genetic data

Previous research suggested that testosterone may influence behavior through dopaminergic pathway [31]. In humans, testosterone administration enhanced activation of the ventral striatum to monetary rewards [32] and the enhancing effects of exogenous testosterone on competitive status-seeking were more pronounced among individuals with a 9/10R compared to 10/10R genotype of the dopamine transporter (DAT) [33]. The expression of DAT, which regulates striatal dopamine, is linked to a 40 base-pair variable number tandem repeat polymorphism of the DAT1 gene [34]. Homozygous 10/10-repeat carriers of this polymorphism have higher DAT expression (i.e., lower striatal dopamine) than heterozygous, 9-repeat variant, individuals [34].

Testosterone’s effects on status-seeking behavior have likewise been shown to be enhanced among individuals with fewer CAG repeats in exon 1 of the androgen-receptor gene [33,36]. In-vitro experimental work suggests that increasing the number of CAG repeats within the androgen receptor (AR) gene reduces the receptor’s transcriptional potential [37]. In other words, the efficiency of the androgen receptors is negatively related to the CAG repeat [38].

We, therefore, tested whether testosterone’s effects on strategic prosociality depended on individual differences in striatal dopamine, assessed by DAT1 polymorphism, and the efficiency of ARs, assessed by the CAG repeat polymorphism.

#### Genotyping of AR CAG repeat and DAT1 polymorphisms

DNA was extracted from buccal swabs and isolated using a resin-based method with Chelex®100 (Sigma Aldrich, USA). For amplification of the CAG repeat polymorphism in exon 1 of the AR gene primers forward - 5’ GCGCGAAGTGATCCAGAAC 3’ tagged with 6–carboxyfluorescein and reverse - 5’ CTCATCCAGGACCAGGTAGC 3’, and for amplification of the DAT1-3’UTR VNTR polymorphism primers forward - 5’ GTCCTTGTGGTGTAGGGAAC 3’ tagged with 6– carboxyfluorescein and reverse - 5’ CTGGAGGTCACGGCTCAAG 3’ were used in PCR with 20 µL reaction using 250 nmol/L final primer molarity. As PCR masterrmix 5x Hot FIREPol Blend Mastermix with 7.5 mM MgCl2 (Solis Biodyne, Tartu, Estonia) was used in all amplifications. The following PCR program was used: initial denaturation step at 95°C for 15 min, followed by 30 cycles each consisting of denaturation at 95°C for 30 s, annealing at 60°C for 30 s and polymerization at 72°C for 1 min. The number of repeats of AR CAG STR and DAT1-3’UTR VNTR was analyzed by fragment analysis using Sanger sequencing on ABI 3500 Genetic Analyzer (Applied Biosystems, USA).

#### Interaction of DAT1 polymorphism with testosterone effects on the correct choice and RLDDM parameters

There were no significant differences in the distribution of the genotype among our experimental groups (χ^2^ (6, *N* = 190) = 5.76 *p* = .451). The 9/10R and the 10/10R genotypes accounted for most of the observed DAT1 genotypes in our sample (36% (N=68) and 56% (N=105), respectively), and we thus used these two genotypes in the analyses by adding DAT1 polymorphisms as a predictor in interaction with the other factors (recipient, drug treatment, visibility) to the GzLMM of correct choice and the GzLMMs of the RLDDM parameters (see *Materials and Methods* in the main manuscript text). The analysis revealed no significant interaction of DAT1 polymorphism with testosterone effect on correct choice (recipient x drug treatment x visibility x DAT1: *OR =* 1.08, *CI* = [0.95, 1.23], *p* = .258), α^posPE^(recipient x drug treatment x visibility x DAT1: *B =* 0.98, *CI* = [0.92, 1.05], *p* = .578), α^negPE^ (recipient x drug treatment x visibility x DAT1: *B* = 1.07, *CI* = [0.98, 1.17], *p* = .141), choice consistency (recipient x drug treatment x visibility x DAT1: *B =* 1.01, *CI* = [1.00, 1.02], *p* = .091) or decision threshold (recipient x drug treatment x visibility x DAT1: *B =* 1.00, *CI* = [1.00, 1.00], *p* = .986).

#### Interaction of AR CAG repeat polymorphism with testosterone effects on the correct choice and RLDDM parameters

Mean-centered CAG repeat lengths of the AR gene in exon 1 were included as a predictor in interaction with the other factors (recipient, drug treatment, visibility) to the GzLMM of correct choice and the GzLMMs of the RLDDM parameters (see *Materials and Methods* in the main text). The analysis revealed no significant interaction of CAG repeat polymorphism with testosterone effect on correct choice (recipient x drug treatment x visibility x CAG: *OR =* 1.00, *CI* = [0.99, 1.02], *p* = .599), α^posPE^ (recipient x drug treatment x visibility x CAG: *B =* 1.00, *CI* = [1.00, 1.01], *p* = .363), αneg^PE^ (recipient x drug treatment x visibility x CAG: *B =* 1.01, *CI* = [0.99, 1.01], *p* = .243), choice consistency (recipient x drug treatment x visibility x CAG: *B =* 1.00, *CI* = [0.99, 1.01], *p* = .576) or decision threshold (recipient x drug treatment x visibility x CAG: *B =* 1.00, *CI* = [1.00, 1.00], *p* = .103).

#### Interaction of trait dominance with testosterone effects on RLDDM parameters

Mean-centered dominance scores [39] were included as a predictor in interaction with the other factors (recipient, drug treatment, visibility) to the GzLMM of correct choice (reported in the main text) and the GzLMMs of the RLDDM parameters. Contrary to the former, the latter analysis revealed no significant interaction of dominance scores with testosterone effect RLDDM parameters: learning rate for positive PE (recipient x drug treatment x visibility x dominance: *B =* 1.01, *CI* = [0.97, 1.04], *p* = .743), learning rate for negative PE (recipient x drug treatment x visibility x dominance: *B =* 1.02, *CI* = [0.98, 1.07], *p* = .331), choice consistency (recipient x drug treatment x visibility x dominance: *B =* 1.00, *CI* = [0.99, 1.02], *p* = .644), or decision threshold (recipient x drug treatment x visibility x dominance: *B =* 1.00, *CI* = [1.00, 1.00], *p* = .578).

### Supplementary information on the questionnaire data

#### Post-task questionnaire

To estimate the subjective perception of being watched, a post-task questionnaire was administered immediately after the end of the reinforcement-learning paradigm. The participants were asked the question: “Did you feel that you were being watched while performing the task?” The answers were classified into three categories: 1 (Not at all), 2 (Moderately), and 3 (Strongly). First, we investigated whether the subjective feelings of being watched were related to the cortisol reactivity to visibility manipulation (see *Manipulation check* in the manuscript’s main text). We next examined whether the subjective feelings of being watched interact with the testosterone administration effect on prosocial choice and RLDDM parameters. As the factors *visibility* and *subjective feeling of being watched* are not orthogonal, we did not add the subjective feelings of being watched as a fourth factor to the analysis. Instead, we replaced the factor *visibility* with the factors *subjective feeling of being watched* and compared whether such a model explains more variance than the original one. We did not find a significant interaction of the factors administration x recipient x subjective feeling of being watched (*OR =* 0.89, *CI* = [0.79, 1.01], *p* = .133). This model did not explain more variance (*Conditional R2*=.102) than the original administration x recipient x visibility model (*Conditional R2*=.103).

#### Portrait Values Questionnaire

We conducted exploratory analyses to examine what motivational constructs may have interacted with testosterone’s effects. To this end, we analyzed data from the Portrait Values Questionnaire [40, 41] which had been administered as part of the study’s extensive questionnaire battery. Note that these are exploratory analyses and that this specific questionnaire was originally intended to be used for analyses related to other tasks that were part of the overall study. They should therefore be considered with caution, and we have clearly labeled them as exploratory and post-hoc. In brief, the Portrait Values Questionnaire is designed to capture the structure of human values explaining the motivational bases of our attitudes and behavior. The questionnaire allowed us to distinguish four principal value orientations: self-transcendence, self-enhancement, conservation, and openness. The four value subscales were separately added to our analysis of recipient x administration x visibility interaction as mean-centered interaction terms. This revealed a significant interaction of *self-enhancement value orientation* with testosterone’s effect on the number of correct choices (recipient x visibility x administration x self-enhancement*: OR* = 0.65, *CI* = [0.48, 0.88], *p* = .006) and interaction of *conservation value orientation* and testosterone’s effects on choice consistency (recipient x visibility x administration x conservation: *B* = 0.60, *CI* = [0.17, 1.02], *p* = .006). The follow-up analyses of these significant interactions are reported in the main manuscript text.

Self-enhancement value did not significantly interact with testosterone effects on choice consistency (recipient x drug treatment x visibility x self-enhancement: *B =* −0.01, *CI* = [-0.06, 0.04], *p* = .633). Conservation did not significantly interact with testosterone effects on correct choice (recipient x drug treatment x visibility x conservation: *OR =* 0.98, *CI* = [0.94, 1.02], *p* = .287). Self-transcendence did not significantly interact with testosterone effects on correct choice (recipient x drug treatment x visibility x self-transcendence: *OR =* 0.97, *CI* = [0.91, 1.03], *p* = .319) or choice consistency (recipient x drug treatment x visibility x self-transcendence: *B =* 0.33, *CI* = [-0.34, 1.00], *p* = .337). Lastly, openness value did not significantly interact with testosterone effects on correct choice (recipient x drug treatment x visibility x openness: *OR =* 1.02, *CI* = [0.98, 1.07], *p* = .328) or choice consistency (recipient x drug treatment x visibility x openness: *B =-*0.01, *CI* = [-0.07, 0.04], *p* = .646).

#### Observers’ salience survey

To estimate the salience of the observers introduced as NGO representatives in our paradigm, we conducted an additional online survey. Using Amazon Mechanical Turk (MTruk) we interviewed N = 73 volunteers (sample size determined by G*Power [42] to achieve 95 % power to detect a small effect size *f* = .20) aged *M* = 34.55 (*SD* = 9.12) years. Participants were asked to perform the task alone and rate how much they agree with the statements regarding different types of observers. Together they rated 7 statements with the following wording, and each statement concerned a different type of observer:

> “As I perform this task on my computer/mobile device alone, I anticipate I would feel differently if………would watch me perform this task.”

The observer’s types were: an unknown other, a friend, a parent, a shop assistant, an NGO representative, a colleague from work, and a technical support worker. A repeated-measures ANOVA comparing the ratings of the respective observers showed that there was a difference across the levels of the within-subject factor *F*(6, 396) = 2.69, *p* = .014, *f* =.204. A follow-up analysis using treatment contrasts and the NGO as a reference level showed a significant difference between the levels of NGO representative and parent (*B* = −0.57, *p* < 0.001), meaning that participants would feel different to a greater extent if they were watched by an NGO representative than a parent. Crucially, no other pairwise comparisons were significant, all *p*s >.169).”

**Figure S1.**
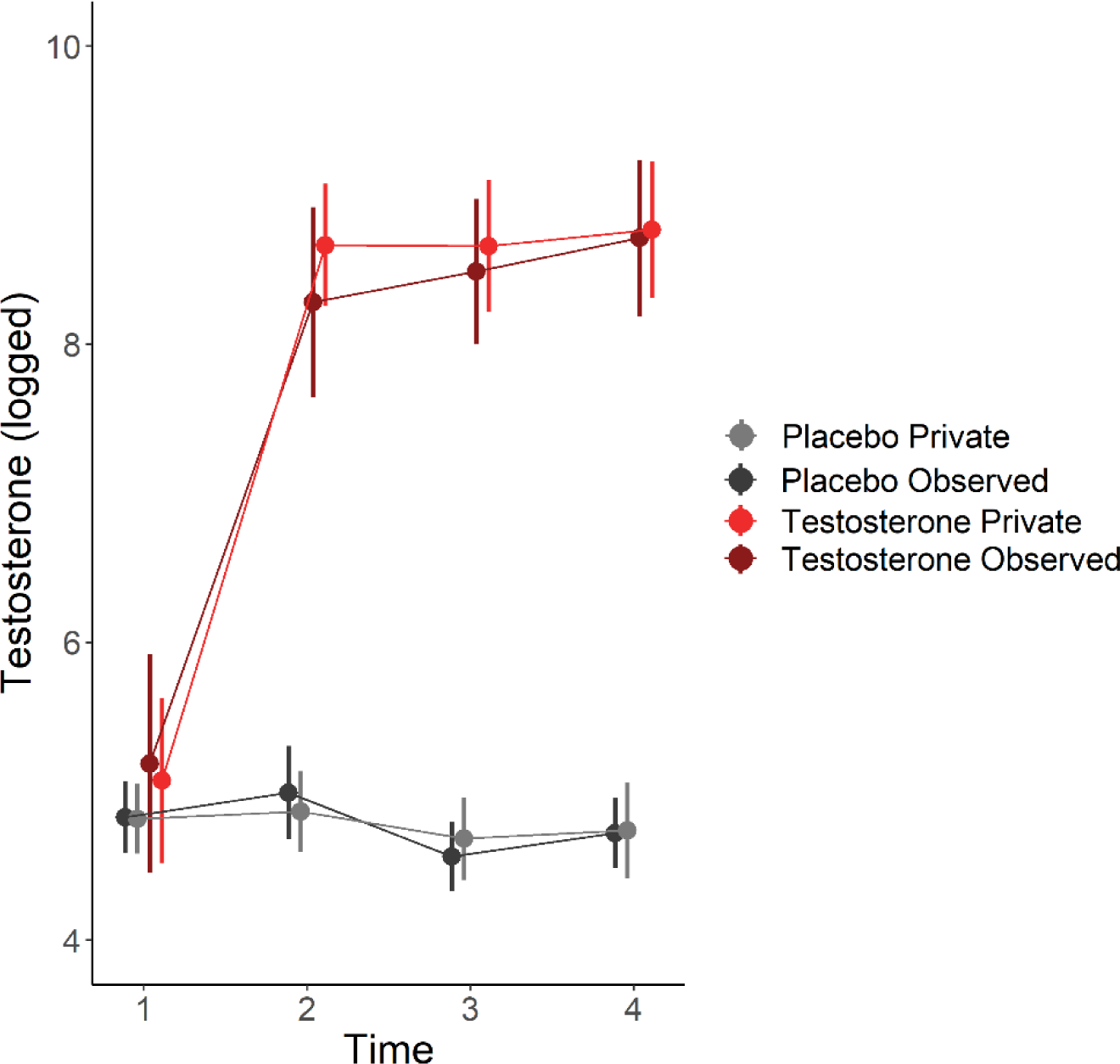
Testosterone levels during the experimental session. Error bars = Mean ± 95%CI.

**Figure S2.**
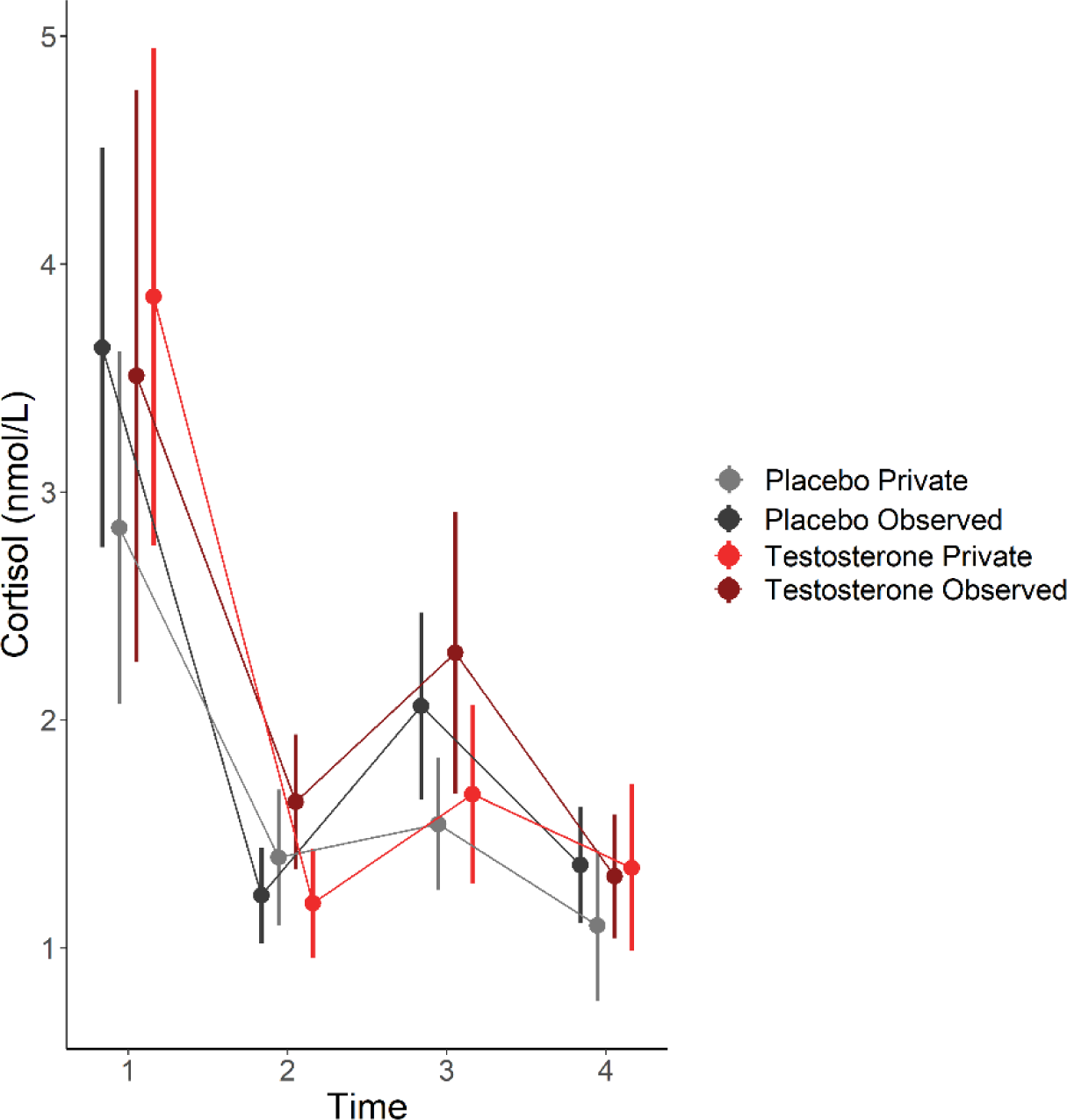
Cortisol levels during the experimental session. Error bars = Mean ± 95%CI.

**Table S1.** Summary statistics across experimental groups: Mean (95%CI), ANOVA.

| Drug treatment | Placebo |  | Testosterone |  | F | p |
| --- | --- | --- | --- | --- | --- | --- |
| Visibility | Private | Observed | Private | Observed |  |  |
| N | 47 | 45 | 53 | 45 |  |  |
| Age | 25.4 (24.5, 26.3) | 24.5 (23.3, 25.7) | 24.0 (23.1, 24.9) | 25.1 (24.0, 26.2) | 1.140 | .239 |
| Baseline testosterone [pg/mL] | 143 (76.7, 209) | 142 (74.9, 209) | 225 (153.5, 296) | 268 (180.9, 354) |  |  |
| log | 4.75 (4.52, 4.98) | 4.76 (4.53, 5.00) | 4.94 (4.69, 5.19) | 4.99 (4.69, 5.30) | 0.024 | .877 |
| Baseline cortisol [nmol/L] | 2.84 (1.81, 3.87) | 3.67 (2.64, 4.71) | 3.89 (2.93, 4.86) | 3.56 (2.52, 4.61) |  |  |
| log | 0.77 (0.52, 1.02) | 0.98 (0.73, 1.24) | 0.90 (0.65, 1.12) | 0.86 (0.60, 1.11) | 0.930 | .336 |
| Baseline estradiol [pg/mL] | 3.68 (3.12, 4.23) | 3.76 (3.20, 4.31) | 3.71 (3.19, 4.22) | 3.75 (3.19, 4.31) |  |  |
| log | 1.23 (1.09, 1.36) | 1.20 (1.06, 1.33) | 1.19 (1.07, 1.32) | 1.23 (1.09, 1.36) | 0.194 | .660 |
| CAG-repeat polymorphism | 19.7 (18.7, 20.7) | 19.8 (18.8, 20.8) | 19.3 (18.4, 20.3) | 19.3 (18.4, 20.5) | 0.001 | .999 |
| Dominance | 3.90 (3.63, 4.17) | 4.02 (3.74, 4.30) | 4.04 (3.78, 4.30) | 3.82 (3.55, 4.10) | 1.526 | .218 |

**Table S2.** Descriptive statistics of cortisol levels during the experiment.

| Drug treatment |  | Placebo |  |  |  |  | Testosterone |  |  |  |  |  |
| --- | --- | --- | --- | --- | --- | --- | --- | --- | --- | --- | --- | --- |
| Visibility | Private |  |  | Observed |  |  | Private |  |  | Observed |  |  |
|  | Mean<br>(CI) | Min | Max | Mean<br>(CI) | Min | Max | Mean<br>(CI) | Min | Max | Mean<br>(CI) | Min | Max |
| Time 1 | 2.84<br>(1.81, 3.87) | 0.64 | 15.34 | 3.67<br>(2.64, 4.71) | 0.37 | 12.87 | 3.89<br>(2.93, 4.86) | 0.23 | 18.59 | 3.56<br>(2.52, 4.61) | 0.44 | 23.80 |
| Time 2 | 1.39<br>(1.13, 1.66) | 0.34 | 5.14 | 1.22<br>(0.96, 1.49) | 0.22 | 3.55 | 1.19<br>(0.95, 1.45) | 0.17 | 4.40 | 1.63<br>(1.33, 1.93) | 0.29 | 3.98 |
| Time 3 | 1.54<br>(1.11, 1.98) | 0.38 | 3.94 | 2.05<br>(1.62, 2.50) | 0.51 | 6.49 | 1.67<br>(1.27, 2.08) | 0.28 | 9.12 | 2.29<br>(1.85, 2.74) | 0.30 | 11.62 |
| Time 4 | 1.09<br>(0.78, 1.42) | 0.21 | 5.73 | 1.36<br>(1.05, 1.67) | 0.47 | 4.86 | 1.31<br>(1.06, 1.64) | 0.15 | 6.57 | 1.35<br>(0.99, 1.63) | 0.15 | 3.77 |

**Table S3.** Model space and model evidence. RW, Rescorla-Wagner model; DLR dual learning rates model; DDM, drift diffusion model; RLDDM, reinforcement learning drift diffusion model; RLDDM-nonlinear, RLDDM with a non-linear transformation function; LOOIC leave-one-out information criterion (lower LOOIC value indicates better out-of-sample predictive accuracy); weight, model weight calculated with Bayesian model averaging using Bayesian bootstrap (higher model weight value indicates a higher probability of the candidate model to have generated the observed data). The winning model is highlighted in bold. Because LOOIC provides relative quantifications of model performance, and the data to be modeled varied across conditions, the absolute magnitude of LOOIC should not be interpreted.

| **Task condition**<br>**Model space** |  | **Placebo Observed** |  | **Placebo Private** |  | **Testosterone observed** |  | **Testosterone Private** |  |
| --- | --- | --- | --- | --- | --- | --- | --- | --- | --- |
|  |  | LOOIC | Weight | LOOIC | Weight | LOOIC | Weight | LOOIC | Weight |
| RW | DDM | 9688 | 0 | 10939 | 0 | 10202 | 0 | 12716 | 0 |
|  | RLDDM | 8365 | 0 | 9747 | 0 | 9135 | 0 | 11450 | 0 |
|  | RLDDM-nonlinear | 8235 | 0 | 9611 | 0 | 8984 | 0 | 11184 | 0 |
| DLR | DDM | 9538 | 0 | 10709 | 0 | 10018 | 0 | 12487 | 0 |
|  | RLDDM | 7877 | 0.004 | 9081 | 0.3 | 8643 | 0.002 | 10688 | 0 |
|  | **RLDDM-nonlinear** | **7832** | **0.996** | **9040** | **0.97** | **8352** | **0.998** | **10516** | **1** |

**Table S4.** Summary of parameters in the winning model. Note that all parameters were further separated for “self” versus “other”, hence for each between-subject condition, the winning model contained 14 parameters. The initial bias *z* in DDM was fixed at 0.5. min(RT), lowest response time from observed data.

| Component | Parameter | Meaning | Interpretation |
| --- | --- | --- | --- |
| RL | $0 < \alpha^{\text{posPE}} < 1$ | learning rate for updating from positive reward prediction error | the learning rate weighs the effect of the reward prediction error in the value update; a higher (lower) learning rate |
| | $0 < \alpha^{\text{negPE}} < 1$ | learning rate for updating from negative reward prediction error | means a faster (slower) value update from the most recent outcome |
| | $\beta > 0$ | inverse temperature in softmax action selection function | choice consistency parameter, captures how much choices rely on the value updates |
| DDM | $v_{\max} > 0$ | maximum value of the drift rate in the non-linear function | defines the upper/lower boundaries in the sigmoid non-linear function |
| | $v_{\text{scaling}} > 0$ | drift scaling that maps value difference into the drift rate | scales the effect of value difference between the choice options on the drift rate |
| | $a > 0$ | decision threshold, i.e., the distance between choice alternatives | “the endpoint” of the evidence accumulation process, captures the amount of information necessary to make a decision |
| | $0 < T < \min(\text{RT})$ | non-decision time | considered to capture sensory delay and/or movement initiation |

